# Multi-omics data integration reveals molecular mechanisms of carfilzomib resistance in multiple myeloma

**DOI:** 10.1101/2024.05.26.595929

**Authors:** Alina Malyutina, Philipp Sergeev, Julia Huber, Juho J. Miettinen, Arnold Bolomsky, Jie Bao, Alun O. Parsons, André Muller, Nara Marella, Mark van Duin, Heinz Ludwig, Jing Tang, Caroline A. Heckman

**Affiliations:** Research Program in Systems Oncology, Faculty of Medicine, University of Helsinki, Helsinki, Finland; Institute for Molecular Medicine Finland (FIMM), Helsinki Institute of Life Sciences (HiLIFE), iCAN Digital Precision Cancer Medicine Flagship, University of Helsinki, Helsinki, Finland; Department of Medicine I, Klinik Ottakring, Wilhelminen Cancer Research Institute, Vienna, Austria; Lymphoid Malignancies Branch, Center for Cancer Research, National Cancer Institute, National Institutes of Health, Bethesda, MD, USA; CeMM Research Center for Molecular Medicine of the Austrian Academy of Sciences, Vienna, Austria; Erasmus University Medical Center Cancer Institute, Rotterdam, The Netherlands

**Author notes:** A.M. and P.S. contributed equally to this study. J.H. and J.J.M. contributed equally to this study.

## Abstract

Multiple myeloma represents a complex hematological malignancy, characterized by its wide array of genetic and clinical events. The introduction of proteasome inhibitors, such as carfilzomib or bortezomib, into the therapeutic landscape has notably enhanced the quality of life and survival rates for patients suffering from this disease. Nonetheless, a significant obstacle in the long-term efficacy of this treatment is the inevitable development of resistance to PIs, posing a substantial challenge in managing the disease effectively. Our study investigates the molecular mechanisms behind carfilzomib resistance by analyzing multi-omics profiles from four multiple myeloma cell lines: AMO-1, KMS-12-PE, RPMI-8226 and OPM-2, together with their carfilzomib-resistant variants. We uncovered a significant downregulation of metabolic pathways linked to strong mitochondrial dysfunction in resistant cells. Further examination of patient samples identified key genes - ABCB1, RICTOR, PACSIN1, KMT2D, WEE1 and GATM - potentially crucial for resistance, guiding us towards promising carfilzomib combination therapies to circumvent resistance mechanisms. The response profiles of tested compounds have led to the identification of a network of gene interactions in resistant cells. We identified two already approved drugs, benidipine and tacrolimus, as potential partners for combination therapy with carfilzomib to counteract resistance. This discovery enhances the clinical significance of our findings.

## Introduction

Multiple myeloma (MM) is a highly heterogeneous hematological malignancy that is characterized by the accumulation of clonal plasma cells in the bone marrow leading to renal dysfunction, anemia, hypercalcemia and bone lesions [1, 2]. It is estimated that there will be 35 780 new MM cases and 12 540 deaths in 2024 in the US [3]. Patients that have been diagnosed with the disease have the median age between 66 to 70 years and the frequency of cases diagnosed before the age of 30 does not exceed 0.3% [4].

The survival of MM patients has been significantly improved over the last fifteen years [5, 6]. Proteasome inhibitor-based therapy approval underlies the advances in both disease outcome and overall survival. Bortezomib and its combinations have been approved for newly diagnosed MM patients, whereas carfilzomib-based therapy is approved for relapsed and refractory cases [6]. Despite steadily improving remission rates, the disease remains incurable with a major hurdle being either intrinsic or acquired resistance to the treatment [7].

Being the first proteasome inhibitor approved for newly diagnosed patients, bortezomib has been extensively studied to reveal molecular mechanisms which underlie its therapy resistance. Genetic abnormalities, atypical metabolic activity, activation of signalling pathways are among the most prominent indications associated with the resistance [8-11]. These hints have led to identification of potential bortezomib combination partners that could overcome the resistance and resensitize the malignant cells to the proteasome inhibitor [12, 13]. At the same time, a second-generation proteasome inhibitor, carfilzomib, has demonstrated its potential to induce cell death in the presence of bortezomib resistance [14, 15]. In contrast to bortezomib, being an irreversible inhibitor, carfilzomib leads to more sustained inhibition of proteasome function. Its high efficacy and selectivity hold great promise for clinical applications which has been supported with positive indication from clinical trials for the newly diagnosed patients [16] (studies NCT01029054, NCT01980589 registered at http://www.clinicaltrials.gov). Despite the explicit advantages, resistance to carfilzomib-based treatment has also been observed to be inevitable for the majority of patients. Several studies have revealed some mechanisms associated with carfilzomib resistance using individual cell line models of resistance[17, 18]; however, rigorous explanation remains elusive.

In this study, we explore the mechanisms of acquired carfilzomib resistance using multi-omics profiling in a panel of four isogenic cell models of carfilzomib resistance (AMO-1, KMS-12-PE, OPM-2, RPMI-8226). Integrated analyses of our basal transcriptome and proteome profiling in matched sensitive and resistant cell variants uncovers an amplification of certain chromosomal regions, elevation of genes/proteins playing role in calcium signalling and potential therapeutic targets that are predicted to have favorable safety characteristics. Our research also identifies metabolic pathway downregulation and mitochondrial dysfunction, validated through various functional assays. Extending the analysis to patient samples from the MMRF CoMMpass trial (version IA15) highlighted six genes: ABCB1, RICTOR, PACSIN1, KMT2D, WEE1 and GATM, contributing to the selection of inhibitors for drug sensitivity tests. Among potential combination partners for carfilzomib, approved drugs benidipine and tacrolimus showed promise, indicating a need for further investigation into their therapeutic efficacy. At the same time, the varied response profiles observed with several tested compounds, alongside their predicted target effects based on chemical structure, have led us to suggest a network of gene interactions as a potential target for drug development.

## Materials and Methods

All methods are reported in supplemental data.

## Results

### Integrated differential analysis revealed elevation of genes from specific chromosomal regions and identified potential targets for novel drug therapies

We conducted differential expression analysis on gene expression from the cell lines, comparing resistant variants to wild type (Fig. 1A). Among 12422 genes analyzed, 2164 met the significance criteria (Table S1). Most of the significant genes were found to be upregulated in resistant cells (N=1152) with ABCB1, NRG4, ZDHHC15, RUNDC3B and SPP1 showing the highest logarithmic fold changes (logFC). A parallel analysis on protein abundance revealed significant differences in 25% of proteins (N=1563 out of 6124, Table S2), with the ABCB1 displaying the most significant logFC, along with other proteins upregulated in resistant cell lines: SRI, MX1, EEF1A2, GBP1 and GBP2.

**Fig. 1.**
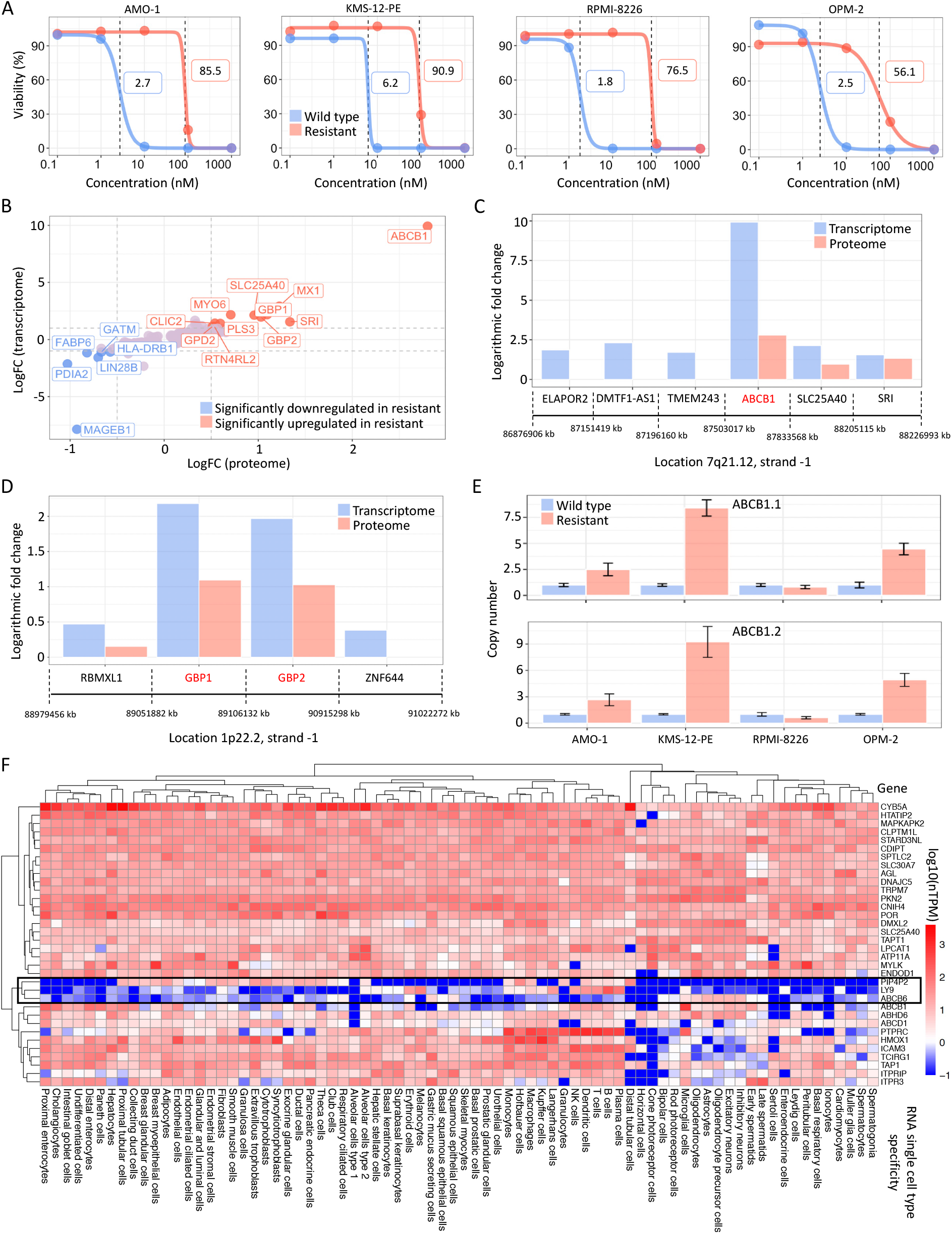
Integrated differential expression analysis key findings. **A** Carfilzomib dose-response curves for wild type and resistant cell lines with IC50 values indicated using dashed lines and numbers. **B** LogFC values for significant genes and proteins. The genes/proteins with the highest absolute logFC are annotated. **C, D** Two chromosomal regions with the highest number of significant genes/proteins and average absolute logFC > 1. The genes’ starting positions on the chromosome are marked with the dashed lines. The last dashed line stands for ending position of the corresponding gene. **E** Copy numbers for the two ABCB1 isoforms in wild type and resistant cell lines. **F** Heatmap displaying log10-transformed nTPM values for 34 upregulated genes/proteins across 79 different cell types.

Combining transcriptome and proteome analysis results narrowed down the list of significant findings to 455, with 444 genes/proteins showing consistent direction in their expression changes. Notably, ABCB1 emerged as the most distinct gene/protein among these findings (Fig. 1B), which is in line with previous reports [17, 19].

We explored further the notable overexpression of ABCB1 in resistant cells and observed that the SRI gene, a close neighbor of ABCB1 on chromosome 7, was also significantly upregulated in these cells (Fig. 1B). Therefore, we extended our analysis to chromosomal regions containing at least 2 significant genes/proteins and considered the band and strand information. This led us to identify two chromosomal regions where the absolute median logFC at both the transcriptome and proteome levels exceeded one (Fig. 1C, D). Indeed, the 7q21.12 region on the negative strand, where ABCB1 is located, contains 6 upregulated significant genes, including 3 significant proteins (ABCB1, SLC25A40 and SRI, Fig. 1C). We confirmed the elevated expression of ABCB1 with Western blotting (Fig. S1) and investigated whether its pronounced upregulation could be attributed to an increase in gene copy numbers. Indeed, in three out of four cell lines examined (AMO-1, KMS-12-PE and OPM-2, Fig. 1E), we identified at least three copies of the ABCB1.1 and ABCB1.2 isoforms. Notably, the KMS-12-PE cell line exhibited the highest copy number, whereas RPMI-8226 mirrored its wild-type variant closely. Another chromosomal region which was identified by our analysis as significant is 1p22.2 region on the negative strand with GBP1 and GBP2 standing out as remarkable genes within this area (Fig. 1D). These genes are part of the larger family of GTPases induced by interferon-gamma. High GBP2 expression correlates with a favourable prognosis in node-negative breast carcinoma patients [20], whereas GBP1 overexpression is linked to a poorer prognosis in oral squamous cell carcinoma cases [21]. At the same time, some studies highlighted a significant GBP-1 role in calcium homeostasis [22, 23].

Yet, GBP1 is not the only gene/protein among the top significant ones that play role in calcium signalling. For example, SRI is a major player of calcium homeostasis - it regulates calcium channels and pumps, endoplasmic reticulum, and cytosolic calcium concentrations. Another protein, MYO6, is a reverse-direction motor protein which binds to calmodulin in its own unique way - through the neck region between motor and IQ domain (Fig. 1B). This interaction necessitates the C-terminal lobe of calmodulin to be bound with calcium, suggesting that a surge in calcium influx may drive MYO6’s overexpression [24]. CLIC2 and GPD2 were also reported to participate in calcium signalling [25, 26].

To identify potential targets with greater clinical relevance, we concentrated on significantly upregulated genes and proteins, investigating their expression across various cell types using data from the Human Protein Atlas (HPA,[27]). We analyzed 34 proteins, employing Euclidean distances for clustering, which we visualized through a heatmap (Fig. 1F). A viable clinical drug candidate should ideally be both effective and safe, meaning it should minimally impact healthy cell populations. Our analysis revealed that the three proteins PIP4P2, LY9, and ABCB6 formed a distinct cluster characterized by considerably lower normalized transcript per million (nTPM) levels across cell types compared to other proteins. Specifically, LY9 exhibited its highest expression mostly in B-cells and plasma cells, with nTPM values of 244 and 102.5, respectively. PIP4P2 showed peak expression in distal tubular cells (nTPM = 49) and Hofbauer cells (nTPM = 48.4). Notably, ABCB6 displayed uniformly low expression in all cell types, remaining below 15 nTPM. Furthermore, we noticed that among the 34 proteins studied, five were plasma membrane proteins, including LY9 (Fig. S2A). Plasma membrane proteins have become a focal point of research due to their accessibility and critical roles in cellular communication and signaling [28, 29], positioning LY9 as a potential therapeutic target. In addition, PIP4P2, LY9, and ABCB6 rank among the top 10 proteins within the HPA annotated group with the highest average expression across both transcriptome and proteome levels, alongside two genes from the ABC family (ABCB1, ABCD1), AGL, MYLK, PTPRC, SLC25A40, and TCIRG1 (Fig. S2B).

These results position LY9 as a promising therapeutic target (eg. for antibody drug conjugates), particularly given its potential for high safety. This assertion is supported by its predominant expression in plasma and B cells, coupled with its significant upregulation at both transcriptome and proteome levels in carfilzomib-resistant cell lines. Such a profile suggests that targeting LY9 could mitigate adverse effects on healthy cells while potentially combating malignant cells.

### Strong mitochondrial impairment and downregulation of metabolic pathways in resistant cells

The integrated analysis highlighted several top downregulated genes/proteins (Fig. 1 B), which were associated with mitochondrial function and metabolism. For instance, GATM, involved in creatine biosynthesis, converts arginine and glycine into guanidinoacetate, the immediate precursor of creatine. Another prominently downregulated gene, FABP6, belonging to the FABP family, participates in fatty acid uptake into cells and the formation of their cytosolic pool [30]. LIN28B plays a pivotal role in regulating cellular metabolism through the LIN28/let-7 pathway. LIN28/LIN28B binding to mRNA of glycolysis and mitochondrial oxidative phosphorylation enzymes promotes their translation [31]. The downregulation of these key players in metabolism prompted a closer examination of metabolic pathways and mitochondrial function in general.

We validated the downregulation of metabolic pathways through gene/protein set enrichment analyses using cell lines. By integrating the transcriptomic and proteomic datasets, we identified 96 significant pathways categorized as metabolic, signalling, or disease by KEGG (Tables S3, S4). Remarkably, most of these pathways exhibited consistent directional changes, with 80 out of 96 pathways showing alignment. Interestingly, a distinct separation in direction was observed when focusing on metabolic and signalling pathways. As depicted in Figure 2A, the upregulated pathways primarily belonged to the signalling group, while nearly all metabolic pathways were downregulated in resistant cells. This observation prompted us to annotate all significant genes/proteins from the integrated differential analysis based on their mitochondrial localization using MitoCarta3.0 [32].

**Fig. 2.**
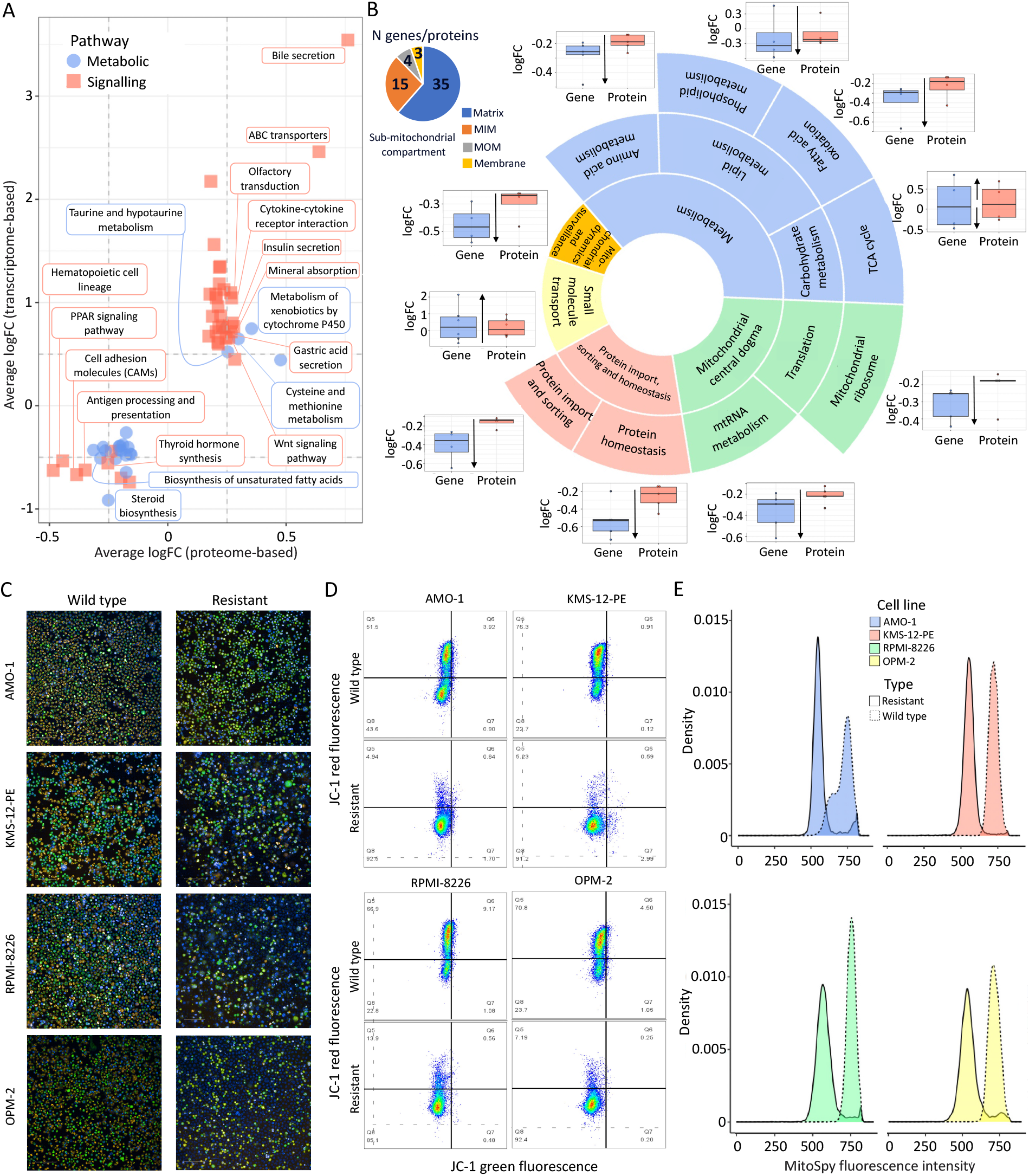
Mitochondrial impairment in carfilzomib-resistant cell lines. **A** Common significant metabolic and signaling pathways from transcriptome and proteome enrichment analyses. Pathways showing the most substantial changes with respect to logarithmic fold change are annotated. **B** Pathways with the highest number of significant mitochondrial proteins, along with their sub pathways are visualized. Distribution of the logarithmic fold changes on gene and protein level for the sub-pathways are captured in the boxplots. The small pie chart indicates the localization of the proteins in the sub-mitochondrial compartments. **C** Representative microscope images and flow cytometry plots (**D**) to evaluate the mitochondrial functional state based on membrane potential of Mitoprobe JC-1 staining in paired wild type and resistant MM cells lines. The presence of red fluorescence signals, indicating the J-aggregated form of the JC-1 dye, marks healthy mitochondria with high membrane potential. Conversely, green fluorescence signifies low membrane potential or depolarized mitochondria. **E** Relative mitochondrial mass differences between wild type and carfilzomib-resistant cell lines based on MitoSpy Green fluorescence intensity.

Through the MitoCarta annotation, we identified 57 proteins, along with their corresponding genes (consistent in direction), localized in mitochondria, predominantly in the matrix (Fig. 2B). The top pathways (N significant proteins/genes > 3) in which these proteins operate are visualized in Figure 2B. Most pathways showed downregulation, except for the small molecule transport, facilitated by ABC family genes and SLC25A40 (located at the ABCB1 amplicon), and the tricarboxylic acid cycle (TCA), which was neutral with half of the genes being upregulated (IDH2 and MDH2) or downregulated (DLD and SUCLG2).

Next, we assessed mitochondrial potential using the MitoProbe™ JC-1 assay kit and detected distinct difference observed between the wild-type and resistant variants through microscope images (Fig. 2C). Quantification of this disparity using the same kit via flow cytometry further highlighted the trend of the resistant cells: they consistently exhibited a notably reduced red fluorescence signal from the J-aggregated form of JC-1 dye, indicative of healthy mitochondria with high membrane potential, resulting in a significant difference in the red to green fluorescence ratio (Fig. 2D, S3). Additionally, we observed relative differences in mitochondrial mass between wild-type and resistant cell lines using the MitoSpy assay. Across all cell lines, there was consistently higher MitoSpy fluorescence intensity in wild-type cells, suggesting a lower mitochondrial content in the resistant variants (Fig. 2E).

Ultimately, we elected to further investigate lysosomal function in the cells, given the critical role of lysosomes in energy production and metabolic balance. We stained the cells with BioTracker™ 560 Orange Lysosome Dye and took the microscopy images. The disparity between the wild-type and resistant variants was not pronounced, with quantitative analysis revealing only a slight reduction in the average lysosomal intensity within the resistant cells (Fig. 3A, B). Additionally, we investigated the relative abundance of autophagic vacuoles. Except for KMS-12-PE cells, all cell lines exhibited higher basal and induced levels of autophagy in wild-type cells (Fig. 3C).

**Fig. 3.**
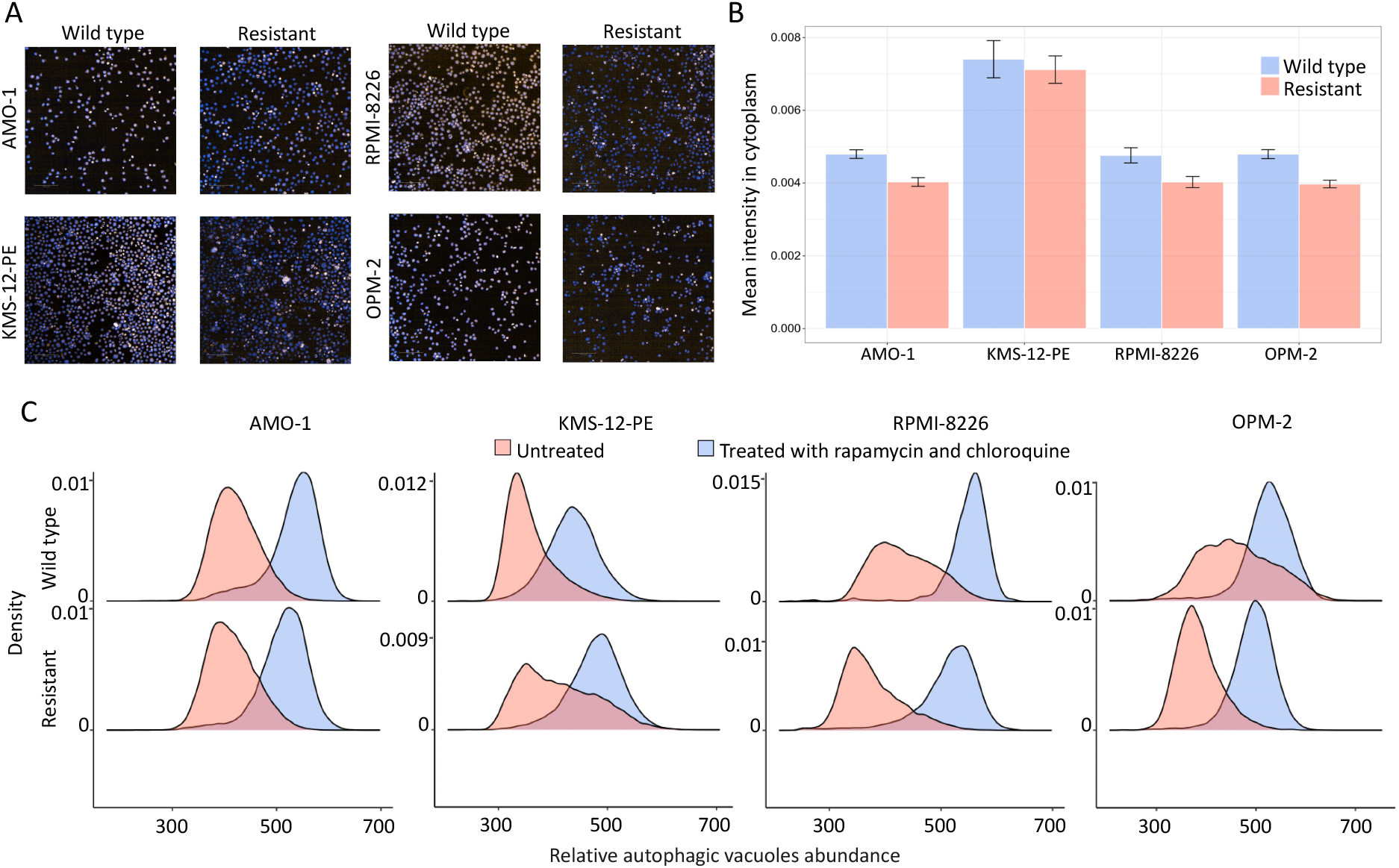
Lysosomal and autophagy assays findings for cell lines. **A** Microscopy images for cell lines stained with BioTracker™ 560 Orange Lysosome Dye. **B** Average lysosomal intensity in cytoplasm in cell lines. **C** Relative abundance of autophagic vacuoles in wild-type and resistant cell lines.

The different effects of carfilzomib on mitochondria and lysosomes might indicate how complex the interactions are between the cell’s recycling and breakdown processes.

### Intrinsic and acquired carfilzomib resistance share common genes and pathways

We next aimed to translate our cell line findings via conducting a differential expression analysis on transcriptomic data obtained from patient samples. Despite contrasting only 6 resistant samples with 41 sensitive ones, we identified 306 significantly different genes, with 255 showing significant upregulation in resistant samples, notably including ABCB1 among the top positive genes (Fig. S4A, Table S5). The enrichment analyses revealed 42 significant pathways, with 19 upregulated in resistant samples (Fig. S4B, Table S6). Intriguingly, all significant metabolic pathways exhibited similar downregulation trends as observed in the cell lines. However, the number of significant genes encoding proteins localized in mitochondria was low (N=13).

Our primary interest was to compare the analyses results between patient samples and cell lines. The overlap between significant genes/proteins in cell lines and patient samples, following similar direction of change, yielded 6 genes: ABCB1, PACSIN1, RICTOR, KMT2D, WEE1, and GATM. Notably, only GATM displayed significant downregulation in resistant cell lines/samples. The difference in ABCB1 expression was more pronounced in both cell lines and samples, while the other genes showed less dramatic changes (Fig. 4A, S4C). Interestingly, a combination of WEE-1 inhibitor, MK-1775, and bortezomib in sequential treatment resulted in more effective treatment than bortezomib alone [13].

**Fig. 4.**
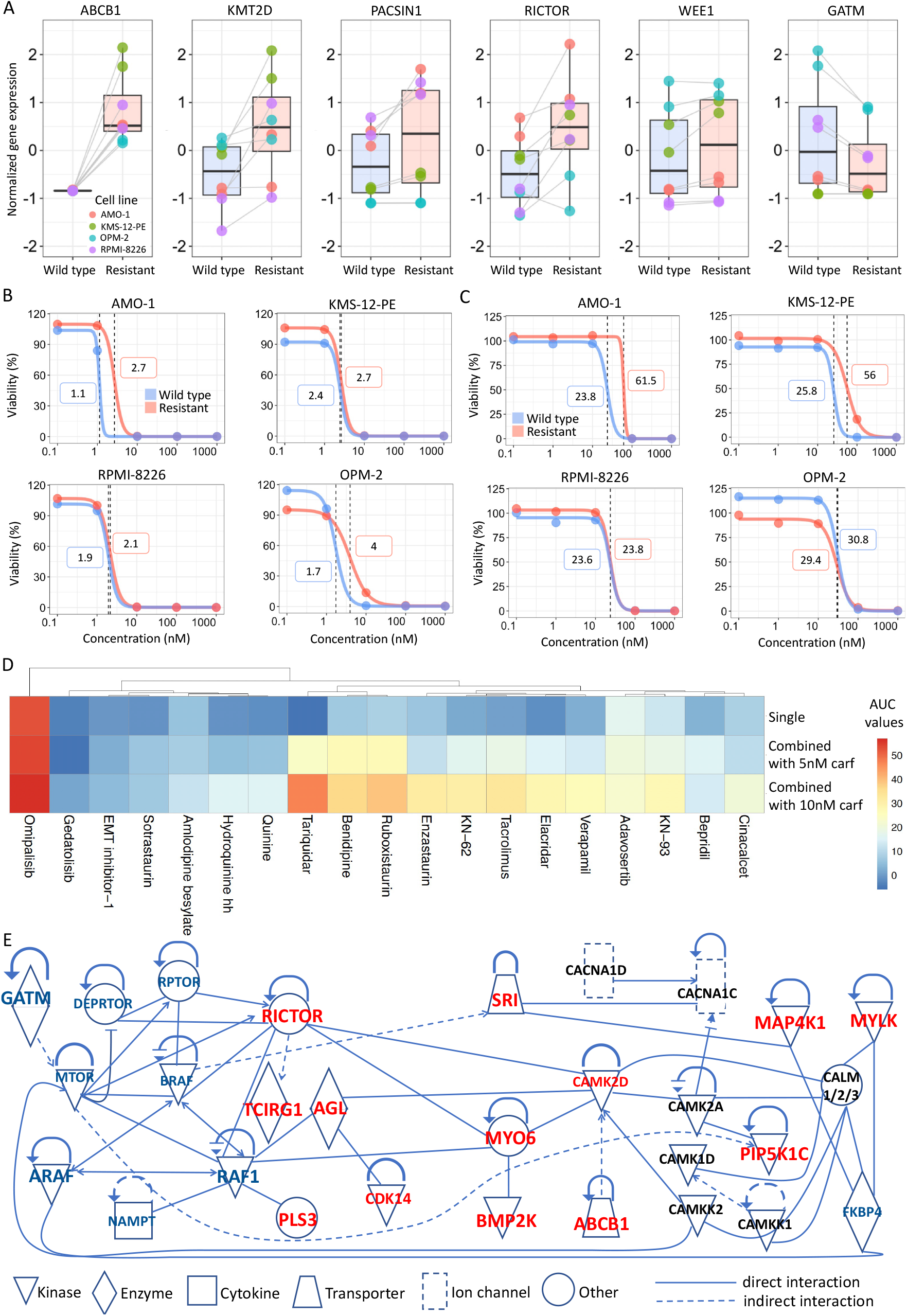
Genes shared by intrinsic and acquired carfilzomib resistance analyses and drug sensitivity assay results. **A** Comparison of the normalized expression levels of common significant genes in wild type and sensitive cell lines. **B** Bortezomib and **C** ixazomib dose-response curves for wild type and resistant cell lines with IC50 values indicated with dashed lines and numbers. **D** Average AUC values for single and combination treatment with either 5 or 10 nM of carfilzomib (carf) in resistant cell lines. **E** Interaction network among significantly different genes identified through computational analysis and some of SEA-predicted drug targets. Genes with larger font sizes are significant at both the transcriptome and proteome levels, with upregulation indicated in red and downregulation in blue. Genes in smaller font are significant in only one type of analysis. Genes marked in black are not significant; instead, they represent predicted targets of the drugs used in the experiments.

Combining the gene/protein set enrichment findings with the patient samples’, we identified 18 pathways consistent in direction (Fig. S4B): one metabolic pathway (carbon metabolism), 9 signalling pathways, and 8 disease pathways. Remarkably, dilated cardiomyopathy and hypertrophic cardiomyopathy were among the shared pathways significantly upregulated in resistant cells suggesting altered expression of genes playing role in calcium signalling. This finding aligns with previous studies, where both events can be triggered by abnormal calcium handling within cardiac muscle cells during carfilzomib treatment [33, 34].

Generally, the limited overlap observed between the cell lines and patient findings could be attributed to the different mechanisms underlying acquired and intrinsic resistance to carfilzomib. Acquired resistance, as anticipated, induces more extensive alterations in the cells. This distinction might reflect how cells adapt over time in response to prolonged exposure to the drug in contrast to intrinsic resistance, which is present from the beginning and might involve different mechanisms.

### Carfilzomib-resistant cells preserve sensitivity to bortezomib and counter resistance with specific combination partners

To counteract carfilzomib resistance, we identified a range of potential combination partners for carfilzomib. Our selection was reasoned by observed increases in the expression of genes/proteins involved in calcium homeostasis (eg. SRI, CLIC2, MYO6, GBP1, GBP2 and GPD2), leading us to incorporate various calcium signalling modulators: calcium channel blockers (such as benidipine, bepridil, and amlodipine besylate), inhibitors of calcium/calmodulin-dependent protein kinase II (KN-93, KN-62), a calcineurin inhibitor (tacrolimus), and a calcimimetic (cinacalcet). While the calcium/calmodulin-dependent protein kinase II inhibitors are not yet approved, the other drugs have been approved for various non-cancer related conditions, like hypertension and angina (Table S7).

In response to significant RICTOR upregulation, observed through both cell line and patient studies, we also added the PI3K/mTOR inhibitors, omipalisib and gedatolisib. Additionally, we included adavosertib, a WEE1 inhibitor, three PKC inhibitors (enzastaurin, sotrastaurin, ruboxistaurin), chosen for the upregulation of PACSIN1, a crucial pathway mediator (Table S7). To target ABCB1 gene, we added first (verapamil, quinine and hydroquinine hydrobromide hydrate) and third generation of ABCB1 inhibitors (tariquidar and elacridar). Prior to inhibitors testing we confirmed upregulation of these four genes in cell lines with Western blotting (Fig. S1). Besides, hippo signalling pathway was the only druggable pathway which was upregulated in cell lines and patient study (Fig. S4B). Due to its potential to inhibit this pathway, EMT-inhibitor was added to the drug panel.

Prior to initiating combination studies, we assessed the efficacy of bortezomib and ixazomib in both wild type and resistant cell lines as a matter of interest. The bortezomib’s effectiveness in resistant cells was remarkably similar to that in sensitive cells, with only minor differences observed (Fig. 4B). Ixazomib showed a similar pattern in RPMI-8226 and OPM-2 cell lines, whereas AMO-1 and KMS-12-PE resistant cell lines displayed a twofold increase in IC50 values compared to their wild type variants (Fig. 4C). Notably, these two cell lines also exhibited the highest resistance to carfilzomib. Despite belonging to the same class of proteasome inhibitors, resistance to one of them does not imply a similar resistance pattern to others.

The combination studies yielded several key findings. Omipalisib stood out for its high efficacy both as a monotherapy and in combination with carfilzomib, achieving average area under the curve (AUC) values ranging from 51.7 to 56.94, suggesting its potential as an effective single agent rather than a combination partner for carfilzomib (Fig. 4D, Table S8). Omipalisib was unique in surpassing the efficacy threshold both alone and when combined with 5nM of carfilzomib. Conversely, gedatolisib, another PI3K/mTOR inhibitor, was among the least effective, regardless of its use as a monotherapy or in combination. This difference in efficacy highlights potential variations in their action specificity and mechanisms despite targeting the same pathway.

At the same time, six additional drugs surpassed the efficacy threshold (AUC>30) in combination with 10nM of carfilzomib: tariquidar (AUC=46.9), ruboxistaurin (AUC=38.8), benidipine (AUC=36.4), tacrolimus (AUC=33.2), enzastaurin (AUC =31.8), and KN-62 (AUC =30.7). Notably, ruboxistaurin and benidipine nearly met the efficacy threshold with 5 nM of carfilzomib, with the corresponding average AUC values being 27.1 and 27.2. Tariquidar, tacrolimus, ruboxistaurin, elacridar, and benidipine emerged as the top five drugs showing the greatest increase in efficacy when used in combination with 10 nM of carfilzomib versus their monotherapy.

While ruboxistaurin displayed high average AUC values in combination with carfilzomib, the other PKC inhibitors, enzastaurin and sotrastaurin, were less effective. Indeed, enzastaurin demonstrated average potency, whereas sotrastaurin showed quite low efficacy (Fig. 4D, S5). Despite targeting the same biological pathway, the distinct mechanisms of action of these drugs could explain their varying efficacies. The secondary targets profiles suggested by SEA for these drugs were extensive, with ruboxistaurin and enzastaurin sharing a greater number of these targets (Fig. S6, Table S9). Sotrastaurin, on the other hand, was categorized separately from the other two, indicating a different range of secondary targets. Similarly, the EMT inhibitor-1, adavosertib, amlodipine besylate, bepridil and remaining ABCB1 inhibitors showed limited efficacy as monotherapies and in combination treatments. Conversely, when combined with carfilzomib, KN-93 and cinacalcet showed modest increases in effectiveness, yet these improvements were insufficient to meet the efficacy threshold.

Driven by varied drug responses within similar classes, we utilized the ingenuity pathway analysis algorithm (IPA, [35]) on key genes/proteins identified in our cell line study along with SEA-predicted drug targets. This approach enabled us to identify a distinct cluster encompassing several major significant genes (Fig. 4E). Majority of the MTOR-interacting genes were significantly downregulated in resistant cells, except for RICTOR. RICTOR acts as a major connector from the downregulated MTOR/BRAF network to the cluster of genes associated with calcium signalling. Although not all genes from the calcium/calmodulin-dependent protein kinase (CAMK) family were marked as significant, they play an essential role in the network. Notably, CAMK2D is a key node, linking RICTOR, ABCB1, MYO6, and AGL with other family members and connecting to MYLK via the calmodulin (CALM) family proteins. Interestingly, MYLK is identified as a predicted target for three drugs—ruboxistaurin, enzastaurin, and benidipine—which are top candidates for combination therapy with carfilzomib based on our drug sensitivity assays. Ruboxistaurin, in particular, displayed the highest Tanimoto coefficient (TC) for MYLK, possibly correlating with its superior efficacy in combination with carfilzomib compared to sotrastaurin, which lacks MYLK as a predictive target and showed a less favourable sensitivity profile. Furthermore, ruboxistaurin is predicted to target all visualized CAMK family members shown in Fig. 4E (Fig. S6). We can hypothesize that CAMK family genes together with MYLK play significant role in survival mechanism of carfilzomib-resistant cells.

## Discussion

MM is a complex blood cancer marked by the proliferation of abnormal plasma cells, leading to various complications and a poor prognosis [1-3]. However, advancements in treatments, particularly with proteasome inhibitors like bortezomib for newly diagnosed patients and carfilzomib as a second-line treatment, have significantly improved patient survival rates [6]. Despite these advances, resistance to carfilzomib is a growing concern, with the underlying mechanisms still not fully understood, indicating a need for continued research in this area.

In this study we addressed carfilzomib resistance via establishing a comprehensive analysis of four carfilzomib-resistant cell lines: AMO-1, KMS-12PE, RPMI-8226, and OPM-2, and their wild type variants. When comparing gene expression in carfilzomib-resistant and wild-type cell lines, a substantial number of genes showed significant changes, predominantly upregulations including the ABCB1 gene (Fig. 1B). Proteomic analysis confirmed these results, with ABCB1 most significantly altered, and a combined analysis of transcriptomic and proteomic data highlighted 455 significant genes/proteins. We have also confirmed ABCB1 amplification (Fig. 1C, E) in three out of four cell lines examined (AMO-1, KMS-12-PE and OPM-2). While ABCB1 amplification have been reported previously to be associated with treatment resistance [36, 37], another amplified chromosomal region which was identified by our analysis as significant, 1p22.2 region on the negative strand with GBP1 and GBP2 genes (Fig. 1D), has not been reported in this context before. However, several studies connect GBP1 overexpression with treatment resistance in cancer [38-40].

ABCB1, also known as MDR1 (multidrug resistance protein 1), encodes an ATP-binding cassette transporter protein that plays a pivotal role in the development of drug resistance in multiple myeloma and other cancers [19, 41, 42]. ABCB1’s function of expelling therapeutic drugs from cancer cells lowers their intracellular levels, reducing drug efficacy and enabling cancer cells to thrive even under treatment. The level of ABCB1 expression in multiple myeloma cells is a critical factor affecting treatment response and patient prognosis [43]. As a key player in drug resistance, ABCB1 has been the focus of therapeutic targeting efforts. Clinical trials have tested ABCB1 inhibitors intending to inhibit its drug efflux capability and enhance treatment effectiveness. However, the clinical advancement of ABCB1 inhibitors has faced challenges, including toxicity and the multifaceted nature of cancer drug resistance mechanisms [44].

In our study, we addressed carfilzomib resistance by looking beyond the direct inhibition of the ABCB1 gene. We explored safety profiles for the significantly elevated genes/proteins in resistant cell lines. PIP4P2, LY9, and ABCB6 were among the top safe proteins based on nTPM expression in various cell types (Fig. 1F). Besides, LY9 is also a plasma membrane protein, which makes him a promising target in monoclonal antibody research field.

Our analysis has also revealed a range of downregulated genes and proteins linked to mitochondrial function and metabolism in resistant cell lines. Enrichment analysis showed a consistent downregulation in metabolic pathways, with an elevation in signalling pathways, suggesting a complex interplay between cellular metabolism, and signalling in resistance mechanisms (Fig. 2A, B). Detailed examination of mitochondrial function revealed that resistant cells exhibited decreased mitochondrial dynamics, including potential and mass (Fig. 2C, D, E). Meanwhile, we observed significant upregulation of key TCA cycle enzymes MDH2, and IDH2 at both transcriptome and proteome levels. This upregulation suggests an increased reliance on the TCA cycle for energy production through the completion of glucose breakdown, generating vital molecules like ATP, NADH, and FADH2 [45, 46]. These findings, combined with RICTOR upregulation on gene and protein level, might indicate a compensatory increase in glycolytic activity to support metabolic needs amidst downregulated metabolic functions and mitochondrial impairment, warranting further investigation.

We extended our study of carfilzomib resistance from cell lines to patient samples (MMRF CoMMpass clinical trial, Version IA15), revealing some consistent findings across both datasets, including a notable upregulation of ABCB1 in resistant samples and metabolic pathways similarly downregulated as in cell line analyses. A direct comparison between the patient and cell line data highlighted five additional genes – PACSIN1, RICTOR, KMT2D, WEE1 and GATM (the only downregulated), which informed the selection of a set of inhibitors for subsequent drug sensitivity testing (Fig. 3A).

In our drug sensitivity testing results, omipalisib notably exceeded the efficacy threshold both as a monotherapy and in combination with carfilzomib, suggesting its potential as an effective single agent. Conversely, gedatolisib showed limited efficacy, underscoring the complexity of targeting the PI3K/mTOR pathway. When assessing the effectiveness of PKC inhibitors in combination with carfilzomib, ruboxistaurin emerged as highly effective, in contrast to enzastaurin and sotrastaurin, which demonstrated reduced potency, particularly sotrastaurin which exhibited the lowest efficacy. While omipalisib and ruboxistaurin are still in clinical trials, approved drugs like benidipine, a vasodilator for hypertension and angina, and tacrolimus, an immunosuppressant preventing organ rejection, emerged as promising carfilzomib partners, meriting further investigation into their combined therapeutic potential (Fig. 3D).

We applied the IPA algorithm to analyze key genes and proteins identified in our cell line studies along with SEA-predicted drug targets. This analysis revealed a distinct cluster of significant genes (Fig. 4E). Notably, while most MTOR-interacting genes were significantly downregulated in resistant cells, RICTOR was an exception, serving as a critical link between the downregulated MTOR/BRAF network and genes involved in calcium signaling.

The CAMK family, though not entirely marked as significant, plays an essential role in this network. CAMK2D links RICTOR, ABCB1, MYO6, and AGL with other family members, and extending to MYLK through CALM family proteins. MYLK is a target for three drugs— ruboxistaurin, enzastaurin, and benidipine—that are top candidates for combination therapy with carfilzomib. Ruboxistaurin stands out, showing the highest TC for MYLK, correlating with its stronger efficacy in combination with carfilzomib compared to sotrastaurin, which lacks MYLK as a target and showed poorer response. Ruboxistaurin is also predicted to interact with all CAMK family members illustrated in Fig. 4E (Fig. S6). Benidipine has both MYLK and ABCB1 among its predicted targets included at the network. Based on the network findings, drug sensitivity assay results and SEA-predicted targets, it is not straightforward to conclude whether targeting ABCB1 leads to higher inhibition of resistant cells than targeting of MYLK or CAMK family members.

A major limitation of this study is the difficulty in obtaining patient samples for studying acquired resistance to carfilzomib, as it’s mainly used in second-line treatment, complicating access to pre- and post-therapy samples. Nonetheless, we managed to find samples where carfilzomib was administered as first-line treatment, aligning some findings with our cell line research. We speculate that exploring samples with acquired carfilzomib resistance might reveal more commonalities, offering further insights into carfilzomib resistance mechanisms. At the same time, despite the limited data, our analysis has generated a substantial number of hypotheses that warrant further exploration. These findings could potentially uncover significant insights and guide future research directions in the field of MM treatment.

## Data availability

The datasets generated during and/or analyzed during the current study are available from the Supplemental Tables. RNA-seq and TMT proteomics data have been deposited to the PRIDE repository. The MM patients clinical and transcriptomic data are available at the MMRF CoMMpass clinical trial data portal (version IA15).

## Acknowledgements

Authors disclose support for the research of this work:

AM: European Research Council (No. 716063), Cancer Foundation Finland, The Päivikki and Sakari Sohlberg foundation and The Instrumentarium Science foundation.

PS: Cancer Foundation Finland

JT: Research Council of Finland (No. 317680), European Research Council (No. 716063).

JB: Research Council of Finland (No. 346019).

CH: Sigrid Jusélius Foundation, Cancer Foundation Finland, Research Council of Finland (No. 334781, 320185, 352265 and 357686).

## Author contributions

AM, PS, JH, JJM, AB, HL, JT and CH conceived the study and designed the experiments, AM analyzed the data, AM wrote the manuscript. PS, JH, JJM, JB, AP, AB, AMe, NM and MvD performed the experiments. All authors reviewed and contributed to the final version of the manuscript.

## Competing interests

CAH: Research funding from Kronos Bio, Novartis, Oncopeptides, WNTResearch, Zentalis Pharmaceuticals for work unrelated to this study, honoraria from Amgen, and personal fees from Autolus.

All other authors have no competing interests to disclose.

