## Supplemental materials for "Multi-omics data integration reveals molecular mechanisms of carfilzomib resistance in multiple myeloma"

### Materials and Methods

#### Patient samples and cell lines

The human multiple myeloma (MM) cell lines used in the study: AMO-1 (ACC 538), KMS-12-PE (ACC 606), OPM-2 (ACC 50), RPMI-8226 (ACC 402) were provided by the German Collection of Microorganisms and Cell Culture GmbH (DSMZ; Braunschweig, Germany). Cells were cultivated according to the DSMZ guidelines and previously described protocols [1] in 1X Roswell Park Memorial Institute (RPMI) 1640 Medium (EuroClone) medium, supplemented with heat-inactivated fetal bovine serum (FBS; Gibco and Lonza), 1% glutamine (Euroclone) and 1% Penicillin-Streptomycin-Glutamine (PenStrep; EuroClone).

Carfilzomib-resistant MM cell variants were obtained after prolonged culture in the presence of gradually increasing concentrations of carfilzomib (PR-171; MedChemExpress) up to a final tolerated dose of 25 nM. We confirmed the carfilzomib resistance via performing drug sensitivity assays on the cell lines to estimate the efficacy of the compound. The carfilzomib-resistant cells showed IC50 values to be at least 14 times higher than their corresponding wild type variants (Fig. 1A).

Short tandem repeat profile analysis was used to test cell lines for authenticity at resistance. All cell lines were tested regularly for mycoplasma contamination using MycoSPY® - PCR mycoplasma test kit (Biontex Laboratories GmbH, Munich, Germany) and used within three months after thawing.

We extracted all clinical and genomic data from the MMRF CoMMpass trial (version IA15, NCT0145429) for MM patients. Since there were no samples available with transcriptomic data both before and after carfilzomib treatment, we focused on intrinsic resistance. Specifically, we included patients who had baseline transcriptomic data and received carfilzomib-based treatment as their initial therapy. Samples were classified as resistant to carfilzomib-based treatment if their therapy response was categorized as 'Stable Disease' or 'Progressive Disease'. Conversely, samples were considered sensitive if the therapy response was marked as 'Stringent Complete Response' or 'Complete Response'. This approach resulted in the selection of 41 sensitive and 6 resistant samples from two treatment groups: carfilzomib + lenalidomide + dexamethasone and carfilzomib + cyclophosphamide + dexamethasone.

#### RNA-seq

RNA of the cell lines was isolated with the AllPrep DNA/RNA Mini Kit (Qiagen). RNA concentration and integrity were determined using a NanoDrop 3300 fluorometer (ThermoFisher Scientific) and a Bioanalyzer 2100 machine (Agilent), respectively. For generating RNA-seq libraries the KAPA mRNA HyperPrep Kit (KAPA biosystems) was used with 250ng RNA input. Paired-end sequencing of libraries was performed on a NovaSeq 6000 (Illumina) machine with 101bp reads.

**TMT-based proteomics**

Cell pellets were lysed in 50mM HEPES (pH 8.0), 2% SDS, 0.1M DTT, 1mM PMSF and protease inhibitor cocktail heated to 99°C for 5 min. DNA was sheared using a Covaris S2 high performance ultrasonicator. Cell debris was removed by centrifugation at 20,000g for 15 min at 20°C and protein concentration determined using BCA. FASP was performed using a 30 kDa molecular weight cutoff centrifugal filters. In brief, 100 µg of total protein per sample was reduced by the addition of DTT to a final concentration of 83.3 mM, followed by incubation at 99°C for 5 minutes. After cooling to room temperature, samples were mixed with 200 µL of freshly prepared 8 M urea in 100 mM Tris-HCl (UA-solution) in the filter unit and centrifuged at 14.000xg for 15 min at 20°C to remove SDS. Residual SDS was washed out by a second wash step with 200 µL UA solution. Proteins were alkylated with 100 µl of 50 mM iodoacetamide in the dark for 30 min at RT. Thereafter, three washes were performed with 100 µl of UA solution, followed by three washes with 100 µl of 50 mM TEAB buffer. Proteolytic digestion is performed using the protease trypsin in a 1:50 ratio overnight at 37°C. Peptides were recovered using 40 µL of 50 mM TEAB buffer followed by 50 µL of 0.5 M NaCl. Peptides were desalted using desalting columns. TMT 7plex Label Reagent Set was used for labeling according to the manufacturer. After the labeling reaction was quenched, the samples were pooled, the organic solvent removed in the vacuum concentrator, and the labeled peptides purified by C18 solid phase extraction. Tryptic peptides were re-buffered in 10 mM ammonium formate buffer pH 10, shortly before separation by reversed phase liquid chromatography at pH 10. Peptides were separated into 96 time-based fractions on a C18 RP column (150 × 2.0 mm, 3 µm) using Dionex Ultimate 3000 series HPLC fitted with a binary pump delivering solvent at 50 µL/min. Acidified fractions were consolidated into 36 fractions via a concatenated strategy. After removal of solvent in a vacuum concentrator, samples were reconstituted in 0.1% TFA prior to LC-MS/MS analysis. Mass spectrometry analysis was performed on an Orbitrap Fusion Lumos Tribrid mass spectrometer coupled to a Dionex Ultimate 3000 RSLCnano system via a Nanospray Flex Ion Source interface.

2D-shotgun LC-MS/MS analysis of whole cell extracts was performed as Data Dependent Acquisition (DDA) on an Orbitrap Fusion Lumos mass spectrometer (ThermoFisher Scientific) coupled to an Dionex Ultimate 3000RSLC nano system (ThermoFisher Scientific) via nanoflex source interface. 1µg was loaded onto a trap column (Pepmap 100, 5µm, 5 × 0.3mm, ThermoFisher Scientific) at 10µL/min using 0.1% TFA. After loading, the trap column was switched in-line with a 50cm, 100µm inner diameter analytical column (packed in-house with ReproSil-Pur 120 C18-AQ, 2.4µm, Dr. Maisch) at 50°C. Mobile-phase A consisted of 0.4% formic acid in water and mobile-phase B of 0.4% formic acid in a mix of 90% acetonitrile and 10% water. 300nL/min and a 210min gradient was used. MS scans were acquired at 375 – 1650 m/z in the Orbitrap at a resolution of 120,000 (at 200m/z of 200). AGC targeted 4x10E5 ions at maximum 50msec. A TopN dependant scan with a cycle time of 3sec set the acquisition of MS2 spectra in the linear ion trap (IT) using rapid scan speed and a quadrupole isolation window of 1.6 Da. HCD was applied with a NCE of 30%. AGC target was set to 1x10E4 with a maximum injection time of 25 msec. Peptide monoisotopic precursor selection (MIPS) was enabled for charge states 2-6 with an intensity threshold set to 5x10E3 and a dynamic exclusion of 90 sec. A single lock mass at m/z 445.120024 was employed [2]. XCalibur version 4.3.73.11 and Tune 3.3.2782.28 were used to operate the instrument.

Following data acquisition, the acquired raw data files were processed using the Proteome Discoverer v.2.4.1.15 platform, with a TMT 7plex quantification method selected.

In the processing step, we used the Sequest HT database search engine and the Percolator validation software node to remove false positives with a false discovery rate (FDR) of 1% at the peptide and protein level under stringent conditions. All MSn spectra were searched against the human proteome (Canonical, reviewed, 20 304 sequences) and appended known contaminants with a maximum of two allowable miscleavage sites. The search was performed with full tryptic digestion. Methionine oxidation (+15.994 Da) and Acetyl (+42.011 Da) Protein N-Terminus were added as Dynamic Modification. Carbamidomethylation (+57.021 Da) of cysteine residues and tandem mass tag (TMT) 7-plex labeling of peptide N termini lysine residues (+229.163 Da) were set as static modifications. Data were searched with mass tolerances of ±10 ppm and ±0.025 Da for the precursor and fragment ions, respectively. Minimum and maximum precursor mass was set to 350 Da and 5000 Da respectively. Results were filtered to include peptide spectrum matches with Sequest HT cross-correlation factor (Xcorr) scores of ≥1 and high peptide confidence assigned by Percolator. MS2 signal-to-noise (S/N) values of TMT reporter ions were used to calculate peptide/protein abundance values. Peptide spectrum matches (PSMs) with precursor co-isolation interference values of > 50, average TMT reporter ion S/N < 10 and SPS Mass Matches < 65 % were excluded from quantification. Both unique and razor peptides were used for TMT quantification. Correction of isotopic impurities was applied. Data were normalized to total peptide abundance to correct for experimental bias and scaled “to all average”. Protein ratios are directly calculated from the grouped protein abundances using an ANOVA hypothesis test. No Imputation was applied. The mass spectrometry proteomics data have been deposited to the ProteomeXchange Consortium via the PRIDE (PubMed ID: 34723319) partner repository with the dataset identifier PXD050215.

**Immunoblotting**

Protein lysates for Western blotting were obtained using complete-lysis-solution (Merck), quantified with Pierce BCA Protein Assay Kit (Thermo Fisher Scientific). Proteins were analyzed with following primary antibodies: GAPDH (#2118; Cell signaling), MDR1/ABCB1 (#12683; Cell signaling), PACSIN1 (ab137390; abcam), RICTOR (#2114; Cell signaling), WEE1 (#13084; Cell signaling) and secondary antibodies: Goat Anti-Rabbit IgH (ab97051; abcam); Anti-mouse IgG (#7076; Cell signaling). Protein bands were detected via Fusion Solo S - EvolutionCapt Solo 6 17,01 imaging system and its applicable FUSION-CAPT software (VILBER).

**Microscopy and image analysis**

For quantitatively analyzing functional mitochondria in each paired cell lines, MitoProbe™ JC-1 Assay Kit for Flow Cytometry (Thermo Fisher Scientific) was used. The experiment was performed on live cells according to the protocol. Briefly, $10^5$ cells were re-suspended in 0.1ml warm completed culture medium (to reach $10^6$ cells/ml concentration). Each cell line was cultured in 4 wells of a 96-well black flat bottom plate (Corning). 10µL of 200µM JC-1(5’,6,6’-tetrachloro-1,1’,3,3’-tetraethylbenzimidazolylcarbocyanine, included in the kit) was added to each well to reach a final 2 µM final concentration. The cells were subsequently incubated at 37°C, 5% CO2, for 30 minutes. Prior to imaging, Hoechst 33342 (Thermo Fisher Scientific) at a final concentration of 1 µM was added to stain the cell nuclei. For live imaging, Opera Phenix high-content screening system (PerkinElmer) was used; 21 images from each well were captured using a 20x objective. To determine mitochondrial quantity, a CellProfiler-based analysis pipeline was established. First, nuclei were segmented using the DAPI channel. Next, cell cytoplasm was identified by propagating 8 pixels from the nuclear boundary. Subsequently, the intensities of the green and red channels were measured in the segmented cell regions of individual cells. Finally, the average measurement of all cells in each image was used for plotting.

For quantitatively analyzing lysosomal functions, the cells were stained with BioTracker™ 560 Orange Lysosome Dye (Sigma-Aldrich) according to the protocol; 25 images from each well were captured using a 20x objective. To determine the lysosomal quantity, the intensity of dye in cell cytoplasmic regions was measured using a CellProfiler, including nucleus and cytoplasm segmentation, and lysosome dye intensity measurement.

**Flow cytometry**

Mitochondrial membrane potential was investigated with membrane-permeant MitoProbe™ JC-1 dye, using protocol recommended by manufacturer. Briefly, 60.000 cells were plated in 96-well plate, and incubated with 100 uM CCCP and 2 uM JC-1 in staining buffer (SB, PBS+0.5% BSA) for 30 minutes at 37 C. After incubation, cells were washed with 100 uL PBS and resuspended in SB, containing 0.1 uM Helix NP NIR reagent for dead cells detection. Novocyte Quanteon flow cytometer was used for data acquisition and analysis. For fluorescence detection, we used V525 and V586 channels.

MitoSpy Green FM (BioLegend) reagent was utilized for measuring mitochondrial mass of individual cells, according to manufacturer's recommendations. In brief, 60.000 cells were plated in 96-well plate, and incubated with the reagent (1 uM) for 30 minutes at 37 C. After incubation we followed the same pipeline as was described earlier. For events detection, B525 channel was used.

Autophagy assay kit (ab139484, Abcam) was used to measure autophagic vacuoles abundance in live cells. Following manufacturer's protocol, cells were plated at the same density as described earlier and were incubated in presence of 0.5 uM rapamycin and 60 uM chloroquine or DMSO overnight at 37C. On the next day, cells were washed, and stained with Green dye (final dilution 1:2000) for 30 minutes at 37C. After incubation, cells were washed again, and the same pipeline for later steps was used as described earlier. Washing was done with assay buffer, included in the kit. For events detection, B525 channel was used.

Flow cytometry results were analysed in FlowJo™ v10.10 Software (BD Life Sciences).

**qPCR**

Cells were lysed in 330 ul TE + 0.1% SDS and 20 µl proteinase K (Thermo Scientific) and incubated at 56 °C for at least three hours. After that they were precipitated with nucleic acid by adding 0.1 volume (35 ul) of three M NaCl and 1 ml of 100% ethanol, mixed and spun down. The liquid phase was removed, and the pellet was resuspended in 300 µl 70% ethanol, cells were spun down and liquid phase was removed. The pellet was then dried for approximately two hours to remove ethanol and resuspended in TE. The final concentration was measured with Qubit BR DNA kit (Invitrogen). The primers used in the experiment can be found in Table S10.

The qPCR was done with HOT-FIREpol EvaGreen qPCR Mix Plus (SolisBiodyne) in a CFX384 (Bio Rad). Primer efficiencies were determined and results analyzed according to the

Pfaffle method using Bio Rad CFX Maestro software. The stability of reference genes was checked from the M-value as determined by the CFX Maestro software.

**Drug sensitivity and secondary targets**

The chemical compounds, DMSO (negative control) and benzethonium chloride (positive control) were preplated to 384-well plates using an acoustic liquid dispensing system Echo 550 (Labcyte). 25-µL cell suspension containing 5.000 cells was added to each drug plate well using a Multiflo FX (BioTek) dispenser. The plates were incubated at 37°C in 5% CO2 for 72 hours. Subsequently, CellTiter-Glo 2.0 (Promega) reagent was added to all wells, and cell viability as luminescence generated by total cellular ATP was measured using a PHERAstar FS (BMG Labtech). The full list of compounds and supplier information can be found in Table S7.

To evaluate a single drug or a combination efficacy, we fit a dose response curve with drda [3] R package and calculated the area under the inhibition curve (AUC) on a logarithmic scale. To make this measure relative we divided it into the area under the maximal possible dose-response curve between the minimal and maximal logarithmic concentrations tested. The normalized AUC values, along with additional parameters from the dose-response analysis, can be found in Table S8. A normalized AUC value of 30 or higher was used as a criterion to indicate the efficacy of a drug or a combination of drugs in resistant cells. This valued was based on the observation that the average AUC for carfilzomib over the range of [0.1, 1000] was 27.9 for resistant cells, while the tested combinations included carfilzomib concentrations up to a maximum of 10 nM.

For some of the drugs, we utilized the Similarity Ensemble Approach (SEA) to predict additional secondary targets based on their chemical structures [4, Table S9]. We applied criteria for considering SEA-predicted targets as reliable, requiring a Z-score greater than 10, a Tanimoto coefficient above 0.4, and a p-value less than 0.05 for human species. Targets not identified by SEA or those failing to meet the specified threshold were assigned a Tanimoto coefficient of zero.

**Computational analyses**

Differential expression analysis (DEA) was applied to the RNA-seq data obtained from cell lines and patient samples. This analysis encompassed two sets of replicates for both wild type and carfilzomib-resistant variants across four cell lines: AMO-1, KMS-12-PE, OPM-2, and RPMI 8226. For DEA, we filtered out low-expressed genes and followed edgeR R package pipeline [5, 6]. We applied quasi-likelihood negative binomial generalized log-linear models to fit the count data and empirical Bayes quasi-likelihood F-tests to obtain gene-specific p- and adjusted p-values [7, 8]. We set the adjusted p-value threshold to be 0.05 allowing for 5% of false positive significant genes. The protein abundances (3 replicates per cell line) were analysed with limma R package using linear model fits followed by empirical Bayes moderation for statistical analysis and similar threshold for significance as in DEA [9]. Gene and protein set enrichment analyses were executed utilizing the EGSEA R package, with a similar adjusted p-value threshold of 0.05 to identify significant pathways [10].

Supplemental figures

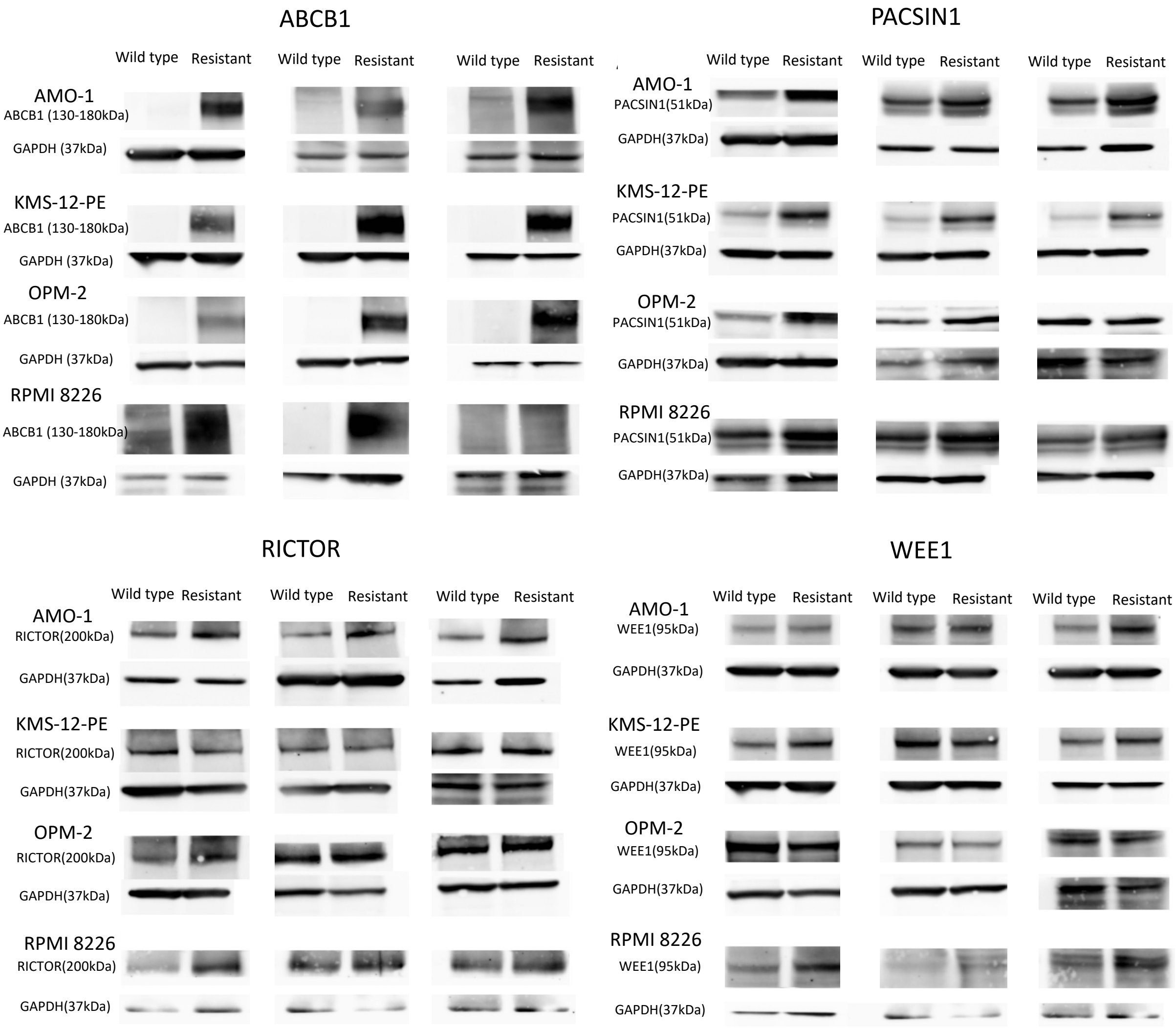


**Fig. S1 Western blots for the 4 druggable genes highlighted by the merged transcriptomic analyses applied to both cell lines and patient samples.**

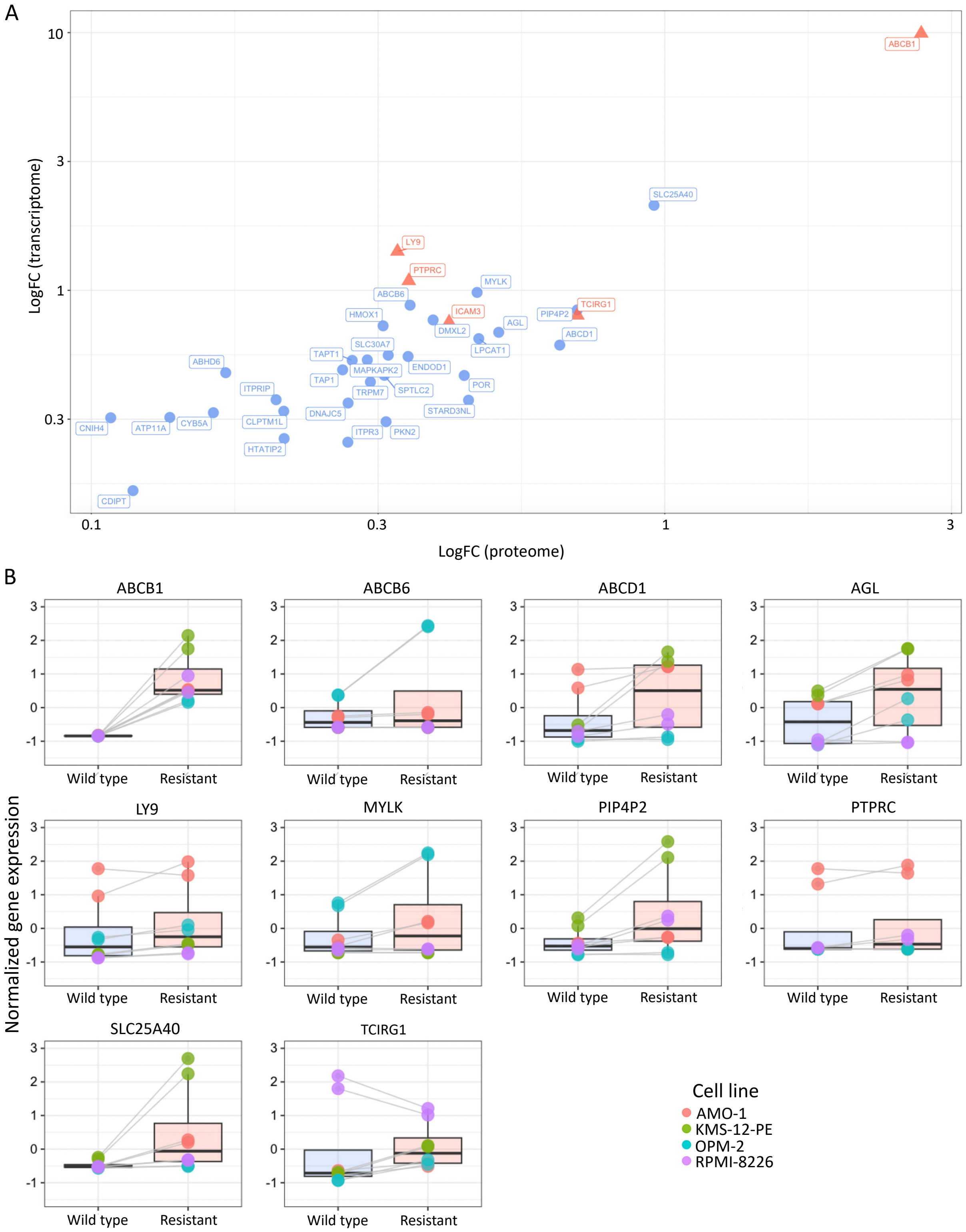


**Fig. S2 Expression levels for genes/proteins that are upregulated in resistant cell lines and annotated for single cell specificity in the Human Protein Atlas (HPA). A** LogFC values measured on transcriptome and proteome levels for the 34 HPA annotated proteins. Proteins located within the plasma membrane group are highlighted in red. **B** Normalized gene expression for the top 10 genes/proteins across the cell lines, ranked based on their average expression at both the protein and gene levels.

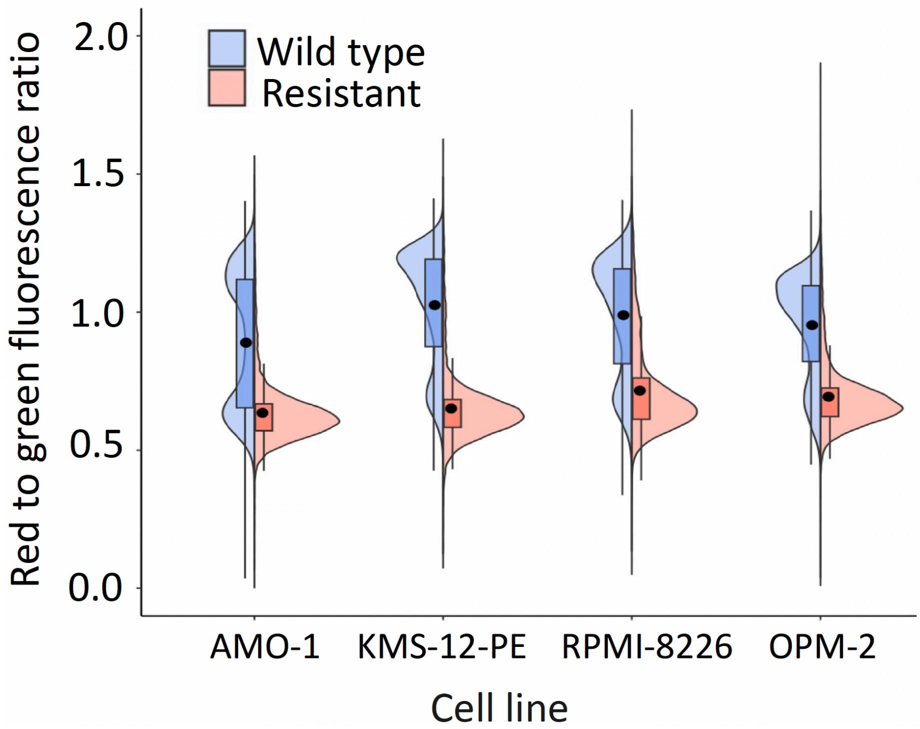


**Fig. S3 JC-1 red to green fluorescence ratio in cell lines.**

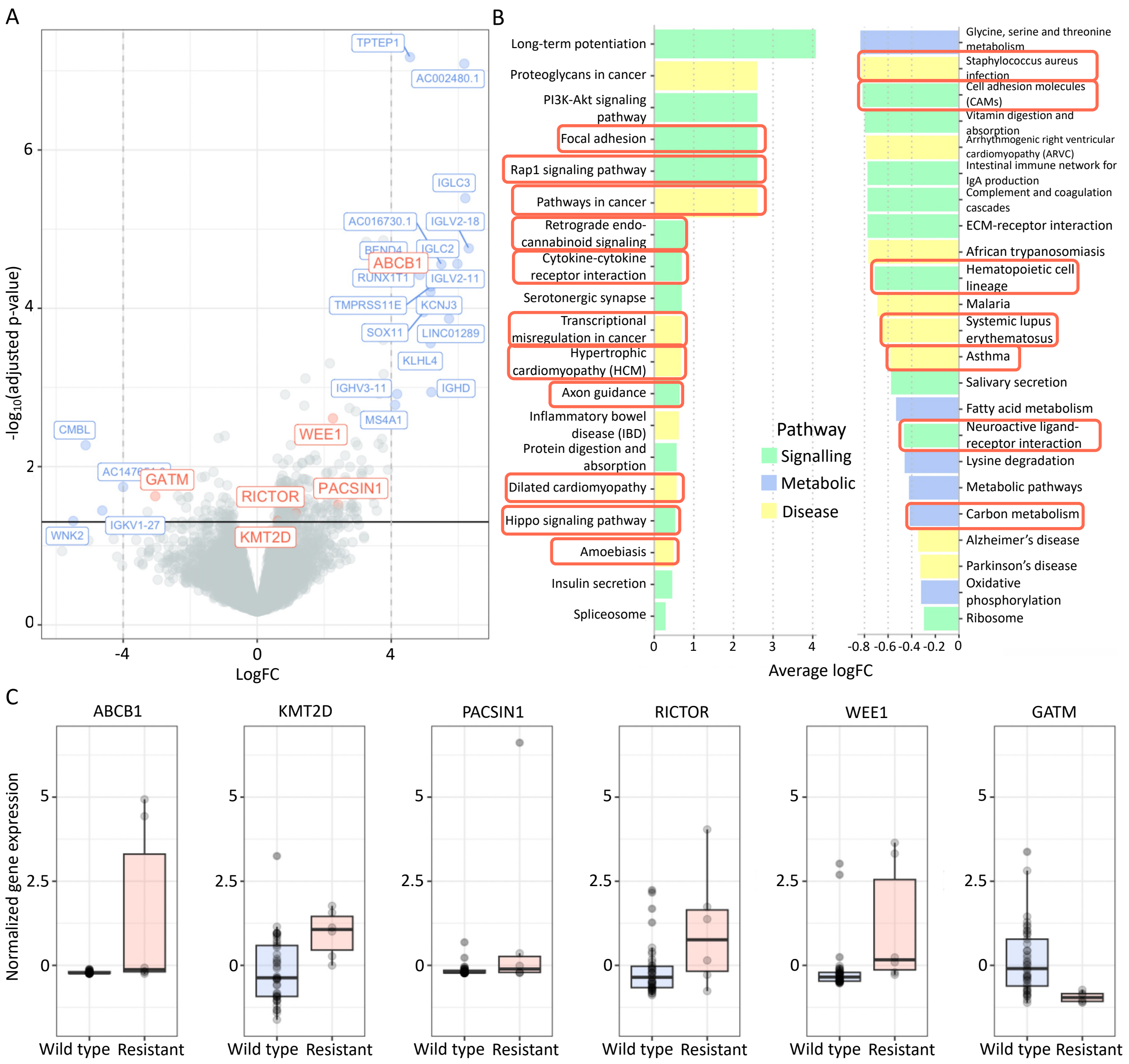


**Fig. S4 Findings from the CoMMpass study. A** Volcano plot displaying genes with the highest logFC, highlighted in blue, and the overlapping genes exhibiting consistent directional changes with the cell line study, highlighted in red. **B** Average logFC for significant pathways resulting from gene set enrichment analyses, with pathways highlighted in red indicating commonality and consistency in direction with the cell line study. **C** Normalized gene expression for the six overlapping significant genes consistent in direction with the cell line study.

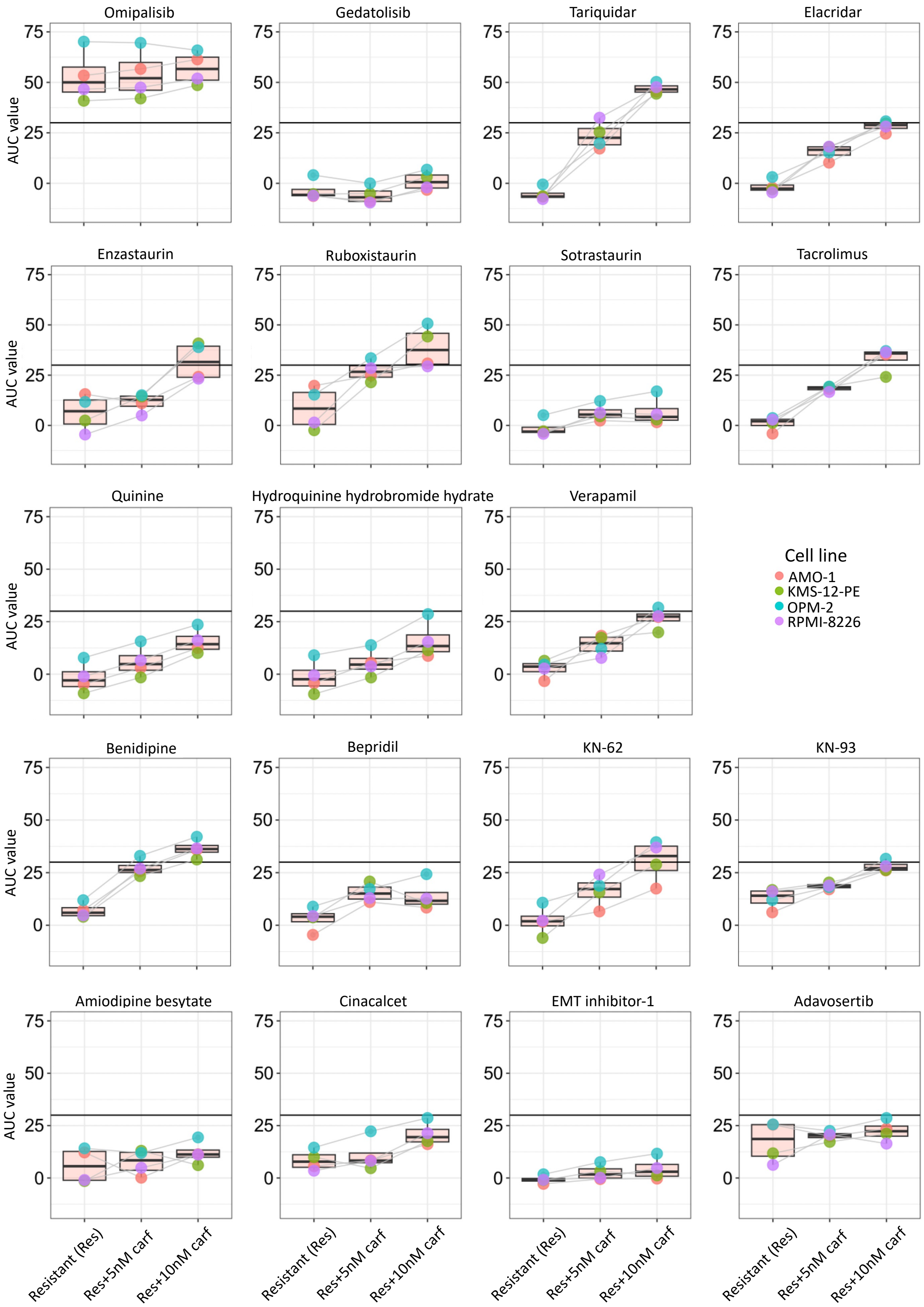


**Fig. S5 Drug sensitivity estimation in resistant cell lines.** The AUC values for dose-response curves are illustrated for single drugs and their combinations with either 5 or 10 nM of carfilzomib (carf). A dashed line indicates the AUC threshold of 30, which was selected to determine a drug's effectiveness.

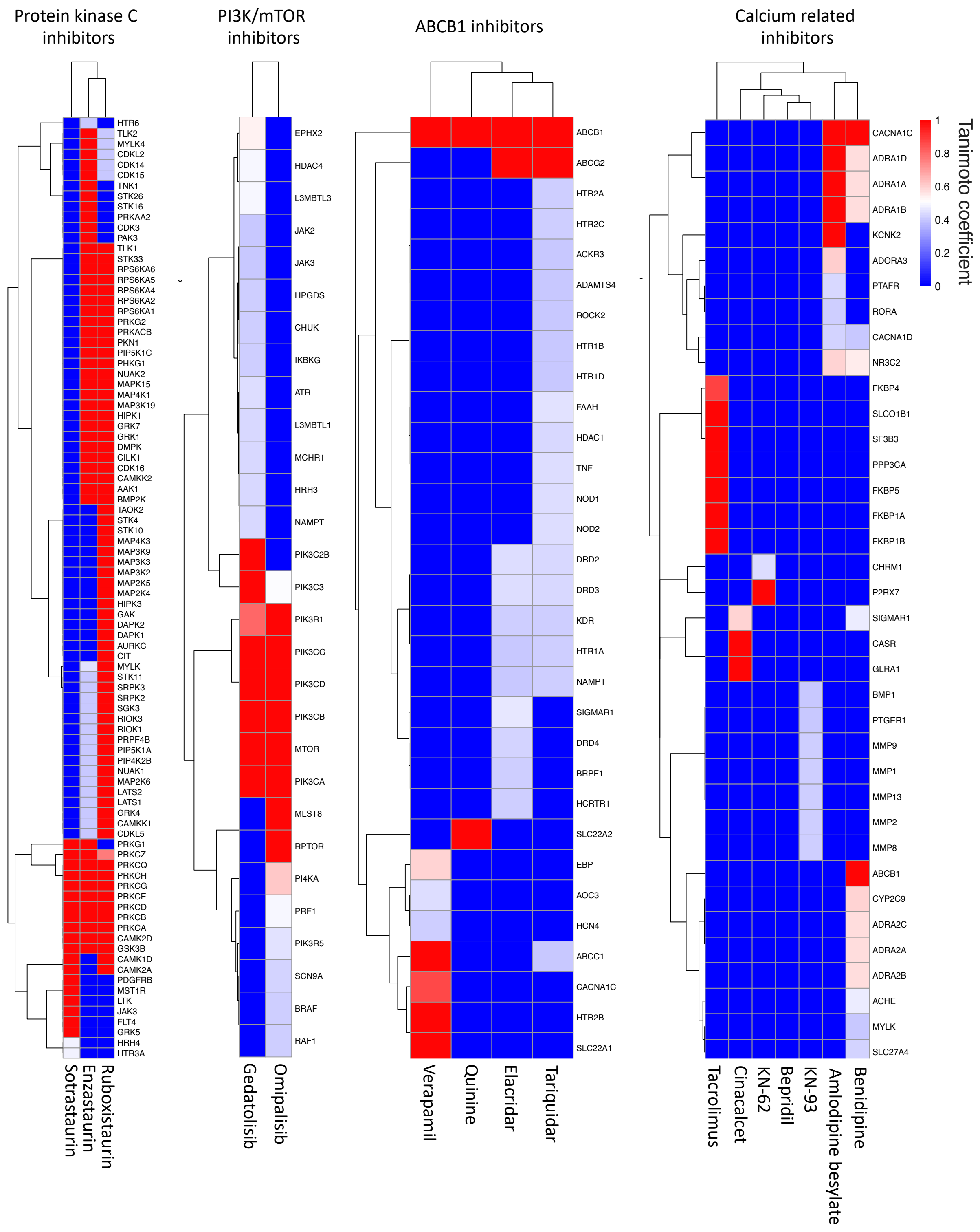


**Fig. S6 SEA-based Tanimoto coefficients for drugs used in the screening. A** Heatmaps with Tanimoto coefficients for the drugs used in the sensitivity assays.
